# PGRP-LA regulates peritrophic matrix synthesis and influences trypanosome infection outcomes in tsetse flies

**DOI:** 10.1101/2025.09.08.674804

**Authors:** Aurélien Vigneron, Brian L. Weiss, Yuebiao Feng, Jingwen Wang, Erick Awuoche, Hang Nguyen, Alessandra Orfano, Liu Yang, Emre Aksoy, Serap Aksoy

**Affiliations:** Department of Epidemiology of Microbial Diseases, Yale School of Public Health, New Haven, CT, USA; Universite Claude Bernard Lyon 1, Laboratoire d’Ecologie Microbienne, UMR CNRS 5557, UMR INRAE 1418, VetAgro Sup, Villeurbanne, France; Fabiotics (Kunshan) Co. Ltd., Jiangsu, China; State Key Laboratory of Genetic Engineering, School of Life Sciences, Fudan University, Shanghai, P. R. China and Ministry of Education Key Laboratory of Contemporary Anthropology, School of Life Sciences, Fudan University, Shanghai, P. R. China; Department of Plant Pathology, Entomology, and Microbiology, Iowa State University, USA; Columbia Center for Translational Immunology, Columbia University Irving Medical Center, New York, NY, USA; Department of Immunology and Infectious Diseases, Harvard T. H. Chan School of Public Health, Boston, MA, USA

**Keywords:** Peptidoglycan Recognition Proteins (PGRP), PGRP-LA, Peritrophic Matrix, vector competence, trypanosome, tsetse fly

## Abstract

Peptidoglycan Recognition Proteins (PGRPs) are conserved pattern-recognition receptors that detect microbe-associated molecular patterns (MAMPs) and activate host immune responses. Compared to other dipterans, the tsetse fly (*Glossina morsitans morsitans*) genome encodes only five PGRPs-PGRP-LA, -LB, -LC, -SA, and -SB – far fewer than most dipterans, likely reflecting its sterile blood diet and streamlined microbiota. Here, we identify PGRP-LA as a critical regulator of peritrophic matrix (PM) integrity in the cardia (proventriculus), the tissue responsible for PM production. The PM is a chitinous sleeve-like barrier that separates the midgut epithelium from the ingested bloodmeal, supporting digestive homeostasis and infection resistance. We show that *pgrp-la* is prominently expressed in the cardia, transiently induced after a bloodmeal in newly eclosed flies, and reinduced following subsequent feedings, likely in response to blood-constituents or mechanical stretch. This induction is sustained during microbial exposure and prolonged in trypanosome-infected flies. RNAi-mediated reduction of *pgrp-la* significantly increased the prevalence of midgut trypanosome infections, indicating a protective role during early infection. PGRP-LA did not mediate infection resistance via canonical IMD pathway signaling, as its silencing did not affect antimicrobial peptide expression. Instead, PGRP-LA modulated the expression of PM-associated genes and gut barrier integrity. Silencing *pgrp-la* reduced PM structure, increased midgut weights and enhanced fly survival following oral challenge with entomopathogen *Serratia marcescens*, likely due to earlier epithelial immune responses through a compromised PM. Similar phenotypes were observed when flies were fed anti-PGRP-LA antibodies, supporting a structural role for PGRP-LA. In addition, soluble variant surface glycoproteins (sVSGs) from trypanosomes and knockdown of *microRNA-275* (miR-275), also decreased *pgrp-la* expression, suggesting that PGRP-LA is part of a broader regulatory network, including the miR-275/Wingless signaling. Collectively, our results identify PGRP-LA as novel regulator of PM biogenesis and vector competence in tsetse, expanding the functional repertoire of PGRPs in insect gut barrier maintenance beyond canonical immune signaling pathways.

**Author Summary:** Insect vectors such as tsetse flies can be infected with pathogens that cause devastating disease in mammals. To protect themselves insect vectors rely on pattern recognition receptors (PRRs) that detect pathogens and activate the production of antimicrobial peptides (AMPs). Physical barriers in the gut also play an important role in limiting infections. One such barrier is the peritrophic matrix (PM), a sleeve-like structure that lines the insect gut and separates the blood meal and its contents from the underlying cells. For trypanosome parasites, which cause sleeping sickness in humans, the PM is the first barrier they must traverse to colonize the tsetse’s gut. In this study, we identified a PRR, PGRP-LA, that, unlike related proteins in other insects that activate AMPs, regulates the integrity of the tsetse’s protective PM barrier. When PGRP-LA was disrupted, the gut barrier weakened, and flies became more susceptible to trypanosome infection. Our work highlights a previously unrecognized role for PGRP-LA in maintaining gut barrier integrity and suggest that targeting this pathway could be a strategy to help reduce parasite transmission.

## Introduction

Tsetse flies (*Glossina* spp.) are vectors of parasitic African trypanosomes, which cause Human and Animal African Trypanosomiases (HAT and AAT, respectively) across sub-Saharan Africa [1]. For successful transmission, these parasites must evade immune defenses in both their vertebrate hosts and invertebrate vectors. In vertebrates, bloodstream form (BSF) trypanosomes evade immune detection through antigenic variation – a process involving the sequential expression of antigenically distinct variant surface glycoproteins (VSGs) on the parasite surface. This strategy allows the parasites to persist despite ongoing host immune responses [2]. In the tsetse fly, both passive and active gut immune barriers eliminate most infections at early stages, restricting transmission to a small fraction of flies [3]. A better understanding of the molecular dialogue between tsetse and trypanosomes, particularly at immune barriers that influence gut colonization success, can inform and facilitate the development of new methods for blocking transmission and reducing disease burden.

Trypanosome infection in tsetse involves a complex developmental process that includes procyclic forms (PF) that colonize the midgut, epimastigote forms (EPF) in the foregut, and mammalian-infective metacyclic forms (MF) in the salivary glands (SG) [4]. The first barrier encountered by ingested BSF trypanosomes is the peritrophic matrix (PM), a chitinous and proteinaceous sleeve-like structure produced primarily by the cardia located at the foregut-midgut juncture. The PM separates the ingested bloodmeal from the gut epithelium, thus regulating the movement of digestive enzymes and protecting the epithelia from harmful compounds in the blood bolus. In this capacity the PM acts as a physical barrier that trypanosomes must circumvent to establish infections [5–7]. Newly eclosed tsetse flies (teneral) have an immature PM that gains functional integrity after the first blood meal. This underdeveloped PM renders teneral flies more susceptible to trypanosome infection than mature adults that have had several blood meals [8–10]. Experimental disruption of PM integrity in older adults, achieved through RNA interference (RNAi) targeting *chitin synthase* or Peritrophin genes that encode PM-associated proteins, significantly increases infection prevalence [11], further supporting the PM’s role as an early barrier to parasite infection.

Similar PM-mediated pathogen resistance functions have been reported in other hematophagous vectors. In mosquitoes, early PM formation restricts *Plasmodium* ookinetes invasion of the midgut epithelium, while delayed PM synthesis enhances parasite transmission [12]. In sand flies, the PM also influences *Leishmania* development by acting as a barrier to parasite migration and modifying the gut environment for parasite survival and development [13, 14]. Thus, the PM has emerged as a critical determinant of vector competence by limiting pathogen establishment during the early stages of infection.

Our previous studies identified parasite-derived factors that influence PM biosynthesis and infection outcome. We found that soluble variant surface glycoproteins (sVSGs), shed into the gut lumen as BSF parasites differentiate into PF forms, are transiently internalized by cardia cells. This uptake leads to downregulation of host *microRNA-275* (*miR-275*), which, by acting through the Wingless/Wnt signaling pathway, reduces expression of several PM protein-encoding genes [15]. This response results in temporary weaking of the PM, which in turn facilitates trypanosome passage across the gut barrier and enhances gut colonization success. The molecular interactions between sVSGs and cardia cells that mediate these downstream effects remains to be elucidated. A similar interaction has been observed in *Anopheles stephensi*, where cell wall components from gut microbiota induce PM formation [16]. The microbiota components were found to act through a pattern recognition receptor (PRR), Peptidoglycan Recognition Protein (PGRP)-LC, which activates the Immune Deficiency (Imd) signaling pathway and induces the expression of the NF-κB transcription factor *Relish* (Rel1), which in turn initiates the expression of the downstream anti-microbial effector proteins. Interestingly, Rel1 also regulates transcription of the gene that encodes the prominent mosquito PM-associated gene, Peritrophin 1 (Per1), suggesting an adaptive link between microbial sensing, immune signaling, and PM biosynthesis [16].

The tsetse fly (*Glossina morsitans morsitans*, *Gmm*) presents a markedly reduced PGRP repertoire compared to closely related dipterans such as *Drosophila melanogaster* and *Musca domestica* [17]. While *Drosophila* [18] and *Musca* [19] encode 13 and 16 PGRP genes, respectively, *Gmm* encodes only five: *PGRP-LC, -LB, -LA,-SA* and *-SB* (named based on their *Drosophila* orthologs [17]). This reduced PGRP repertoire is thought to result from the highly streamlined microbiota that tsetse houses, as a consequence of its strict vertebrate blood diet and viviparous reproductive strategy during which all immature stages develop within the maternal uterus [20]. Among these PRRs, *Gmm* PGRP-LC activates the Imd pathway, which induces the synthesis of antimicrobial peptides (AMP) [21, 22], including Attacin, that exhibit trypanocidal activity [23]. PGRP-LB, in contrast, is highly expressed in the fly’s bacteriome where it promotes mutualism with tsetse’s obligate endosymbiont *Wigglesworthia* by degrading PGN that could lead to harmful immune activation [22]. While the amidase function of PGRP-LB benefits host reproductive fitness [24], secreted PGRP-LB also exhibits direct trypanocidal activity and limits parasite infections in the gut [24, 25].

Here we investigated the role of *Gmm* PGRP-LA, which is prominently expressed in gut epithelia of other insects. In *Drosophila,* PGRP-LA regulates epithelial immune responses during bacterial infection [26], while in *Anopheles* mosquitoes, reduced of *pgrp-la* expression is associated with decreased levels of gut immune effectors that regulate microbiota density and *Plasmodium berghei* infection success [27]. To assess its function in tsetse, we characterized *pgrp-la* expression across distinct gut compartments during the blood feeding process, as well as temporally following immune-challenge and in fly’s that house mature trypanosome infections. Using functional genomic approaches, we examined the role of PGRP-LA in immune regulation, its potential interaction with PGRP-LC mediated Imd-signaling pathway, and its influence on gut PM barrier integrity. We discuss our findings in the context of the non-canonical role insect PGRPs serve in gut homeostasis and structural barrier function, with important implications for vector competence.

## Results

### *Glossina* PGRPs that are responsive to trypanosome infection

The spatial distribution and immune-responsive profiles of *pgrp-lb* and -lc in response to tryp infection have been previously characterized [22, 28]. However, little is known the spatial expression of *pgrp-la* and -sc and how they respond to trypanosome infection. To address this gap, we first measured changes in transcript abundance of *pgrp-la* and -*sa* to trypanosome infections in different gut compartments. Specifically, we performed qRT-PCR to quantify their expression across three gut regions, cardia, bacteriome, and midgut, in age-matched uninfected and trypanosome [*Trypanosoma brucei brucei* (*Tbb*) strain RUMP 503] infected adult flies. While *pgrp-sa* expression showed only a modest increase in gut tissues of trypanosome infected flies (Figure 1A, p=0.05), *pgrp-la* expression significantly was significantly higher in the cardia, bacteriome, and midgut of parasitized individuals when compared to uninfected controls (Figure 1B). The significant increase in *pgrp-la* expression, especially in the cardia, suggests that PGRP-LA may play an active role in modulating the tsetse immune response(s) to trypanosome infections, which we further investigated.

**Figure 1.**
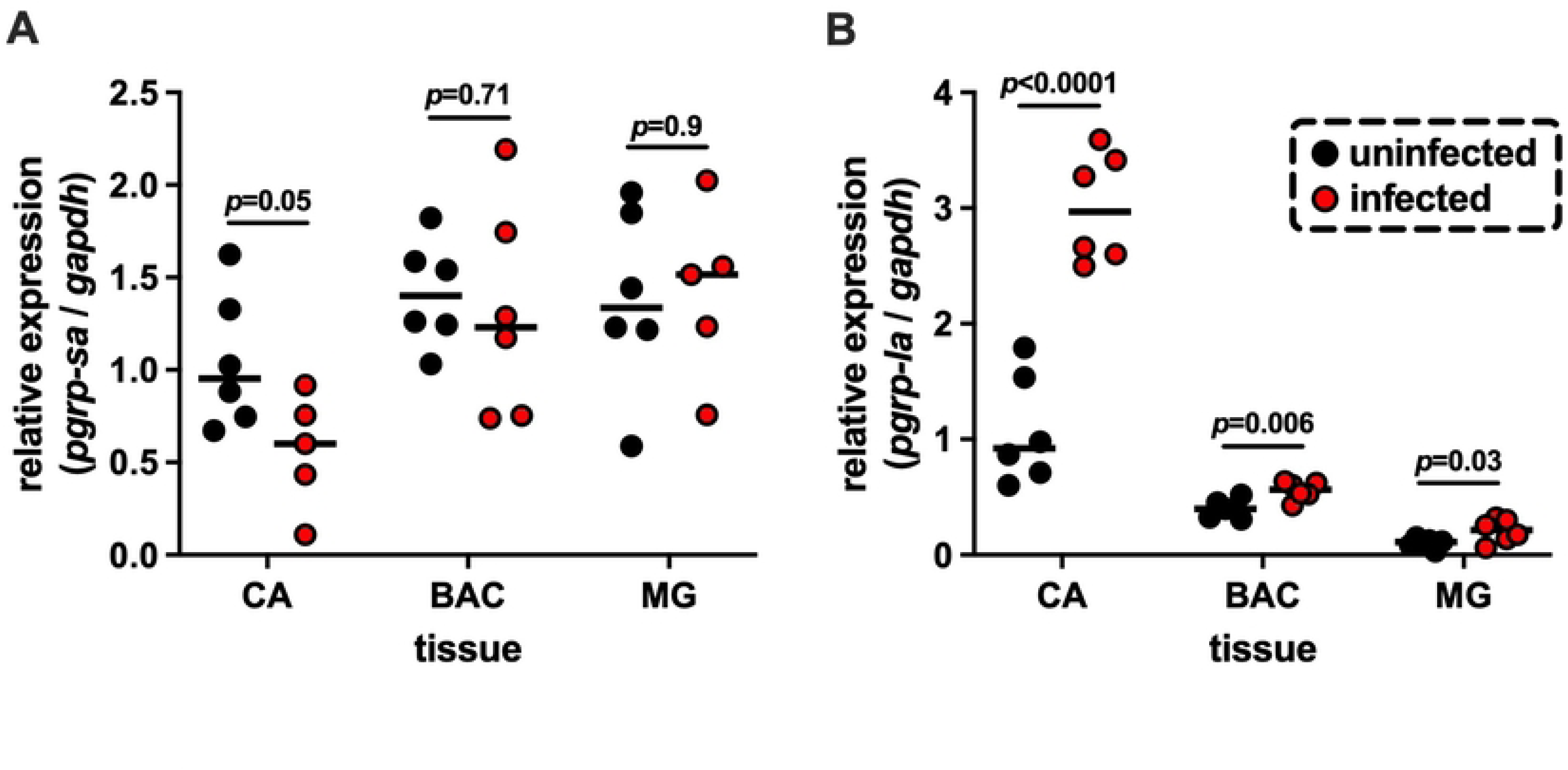
Expression profiles of tsetse *pgrp-sa* and *pgrp-la* in different gut compartments in response to trypanosome infection. Relative expression of (A) *pgrp-sa* and (B) *pgrp-la in* the cardia (CA), bacteriome (BAC), and midgut (MG) of uninfected flies and 14-day old *Trypanosoma brucei brucei* infected flies. Each data point represents an individual biological replicate, containing of pooled tissues from five flies. Black circles indicate uninfected controls and red circles indicate infected tissue samples. Median values are shown as horizontal black bars. Gene expression was measured by qRT-PCR and normalized to the expression of the housekeeping gene *gapdh*. Statistical significance was determined by two-way ANOVA followed by Tukey’s multiple comparisons test (GraphPad Prism v.10.4.1).

### *Glossina* PGRP-LA exhibits high intra-genus conservation

The *pgrp-la* locus resides adjacent to the *pgrp-lc* coding region in the six *Glossina* species for which whole genome sequence (WGS) data is available, except for *G. brevipalpis*, where this locus is not annotated [17, 29]. We first compared the structure of *Gmm pgrp-la* (coding sequence) with that of its orthologs in other Diptera. The putative *Gmm* PGRP-LA encodes a transmembrane protein with a N-terminal cytoplasmic RIP Homotopic Interaction Motif (RHIM)-like domain that is 95% identical in amino acid composition to its *Drosophila* ortholog (Figure 2A, S1A). In *Drosophila*, the RHIM domain of PGRP-LC is necessary for activating the IMD pathway [30]. However, the RHIM domains of *Drosophila* and *Glossina* PGRP-LAs differ from that of PGRP-LC in that they contain a Valine and Isoleucine substitution (Figure S1B) [30]. Although these amino acid residues do not appear to be critical for the interaction of *Drosophila* PGRP-LA with the IMD pathway [26], it remains to be seen whether tsetse PGRP-LA also interacts with the IMD pathway.

**Figure 2.**
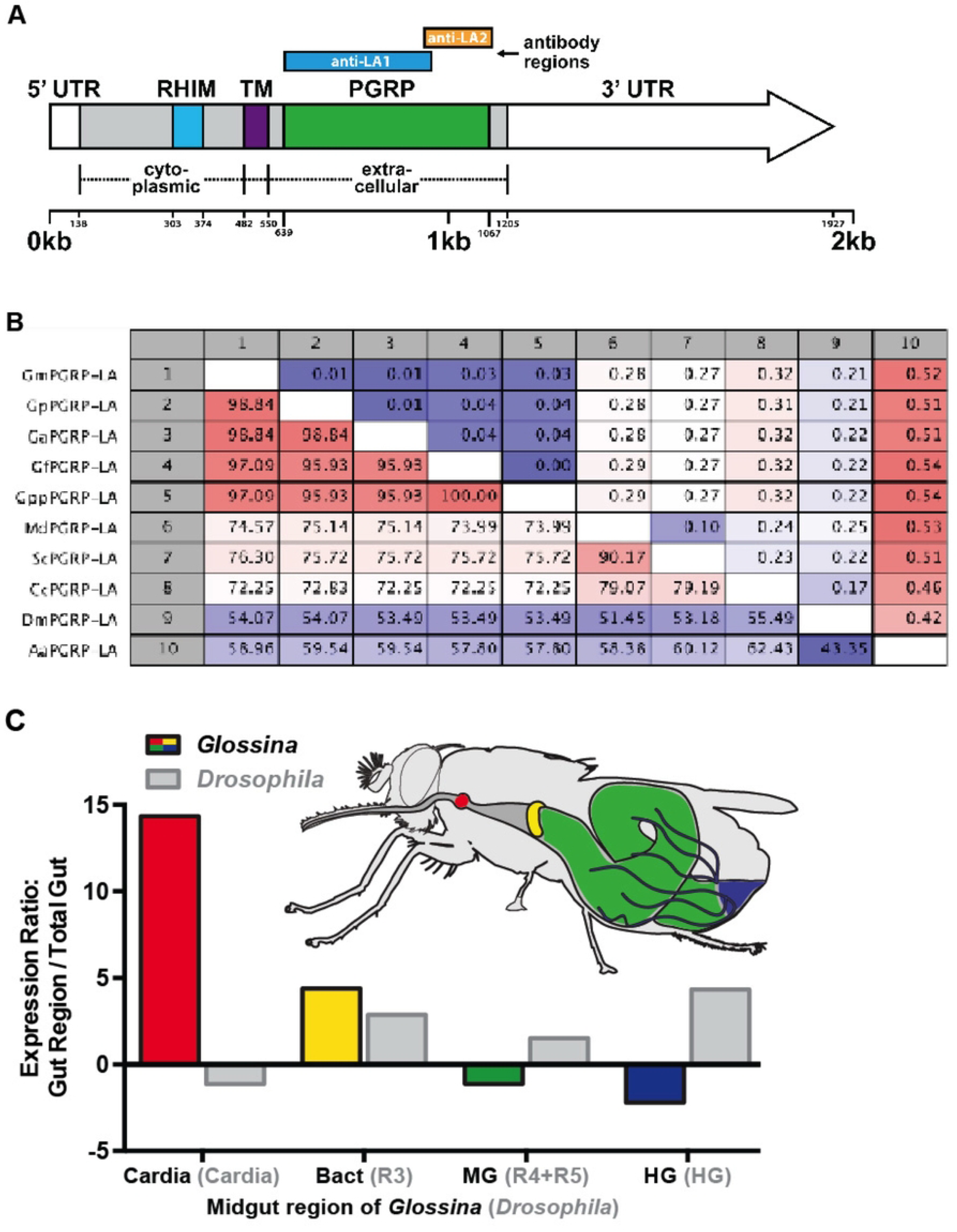
Structure and spatial expression profile of *pgrp-la* in tsetse fly midgut compartments. (A) Schematic representation of the *pgrp-la* locus and its putative product. The predicted PGRP-LA protein includes an extracellular PGRP domain (green), a transmembrane domain (TM, dark purple) and a cytoplasmic RHIM-like domain (blue). No signal peptide was predicted. Coding sequence (CDS) regions lacking known functional domains are shown in gray. The regions used to generate the recombinant proteins for antibody production (anti-LA1 and anti-LA2) are also indicated. (B) Distance matrix showing orthology of the predicted *pgrp-la* domain across fly species. Species abbreviations are as follows: Gm (*Glossina morsitans*), Gp (*Glossina pallidipes)*, Ga (*Glossina austeni*), Gf (*Glossina fuscipes*), Gpp (*Glossina palpalis palpalis)*, Md (*Musca domestica)*, Sc (*Stomoxys calcitrans)*, Cc (*Ceratitis capitata*), Dm (*Drosophila melanogaster*), and Aa (*Aedes aegypti)*. (C) Schematic of the tsetse fly digestive system showing the regions analyzed for *pgrp-la* expression. The cardia (red) marks the beginning of the midgut (MG, green) and delineates this tissue from the foregut (not shown). The bacteriome (Bact, yellow) harbors the obligate intracellular endosymbiont *Wigglesworthia*. The Malphigian tubules (black, wavy lines) connect to the gut at the junction between the midgut and hindgut (HG, blue). Expression of *pgrp-la* was analyzed in each tissue by qRT-PCR using 6 biological replicates per tissue with each replicate containing pooled tissues form five individuals. Expression of *pgrp-la* was normalized to the housekeeping gene *gapdh*. For gut regions, *pgrp-la* expression was normalized to that of its *expression* in the whole gut (*n*=12 biological replicates, each from five pooled individuals). For comparison, *Drosophila* midgut expression data were obtained from the FlyGut atlas, and the corresponding *Drosophila* gut regions indicated are indicated in gray parentheses on the X-axis.

In many PGRPs, the PGRP domain mediates peptidoglycan (PGN) binding. Distance Matrix analysis shows that the putative PGRP domain of tsetse PGRP-LA is highly conserved across *Glossina* species, with amino acid sequence identities ranging from 95.93% to 100% (Figure 2B). In comparison, sequence identity between tsetse PGRP-LAs and those from other higher Dipterans ranges from 72.25% to 76.30%, and between 54% to 60% when compared to *Aedes aegypti*. Over the 172 amino acid residues that constitute the PGRP domain, 25 are unique to tsetse PGRP-LA (Figure S1C). Interestingly, comparative analysis of residues critical for PGN interaction in *Drosophila* PGRP-LCx indicates that only four of the 25 residues are retained in tsetse PGRP-LA (Figure S1C). It remains to be seen whether tsetse PGRP-LA can bind PGN or whether it might serve other regulatory roles such as a negative regulator or co-receptor rather than a primary sensor.

### *pgrp-la* is preferentially expressed in tsetse’s cardia and bacteriome tissues

We evaluated the spatial expression profile of *pgrp-la* across four compartments of tsetse’s gut: the cardia, bacteriome, midgut, and hindgut (Figure 2C). We also quantified the overall expression of *pgrp-la* in the entire gut (from foregut to hindgut) to calculate fold-changes of expression within specific compartments relative to the whole gut. We observed that *pgrp-la* transcript abundance is approximately 15 times higher in the cardia and 5 times higher in the bacteriome compared to the whole gut (Figure 2C). In contrast, *pgrp-la* transcript levels in the midgut and hindgut are about 1.2 and 2 times lower, respectively, compared to the whole gut. This expression pattern differs from that observed in *Drosophila*, where *pgrp-la* is expressed four times higher in the hindgut than in the whole gut, with no significant difference noted in the cardia according to data presented in FlyBase (Figure 2C). These spatial variations in *pgrp-la* expression further support our theory that the function of the encoded protein may be different in *Glossina* than in *Drosophila*.

### *pgrp-la* expression in response to a blood meal and microbial challenge

We next examined *pgrp-la* expression in the cardia of teneral flies, and at 6, 24 and 72 hours (hrs) following their first bloodmeal, as well as 6 and 24 hrs after a second bloodmeal given to flies starved for 72 hrs (denoted 72-6 and 72-24, respectively). Our results indicate that *pgrp-la* expression was significantly upregulated at 6 hrs post bloodmeal and again at 72-6 hrs following a second bloodmeal. However, expression returned to baseline by 24 and 72-24 hrs (Figure 3A), suggesting that *pgrp-la* is transiently responsive to blood ingestion, possibly via mechanical or nutritional signals.

**Figure 3.**
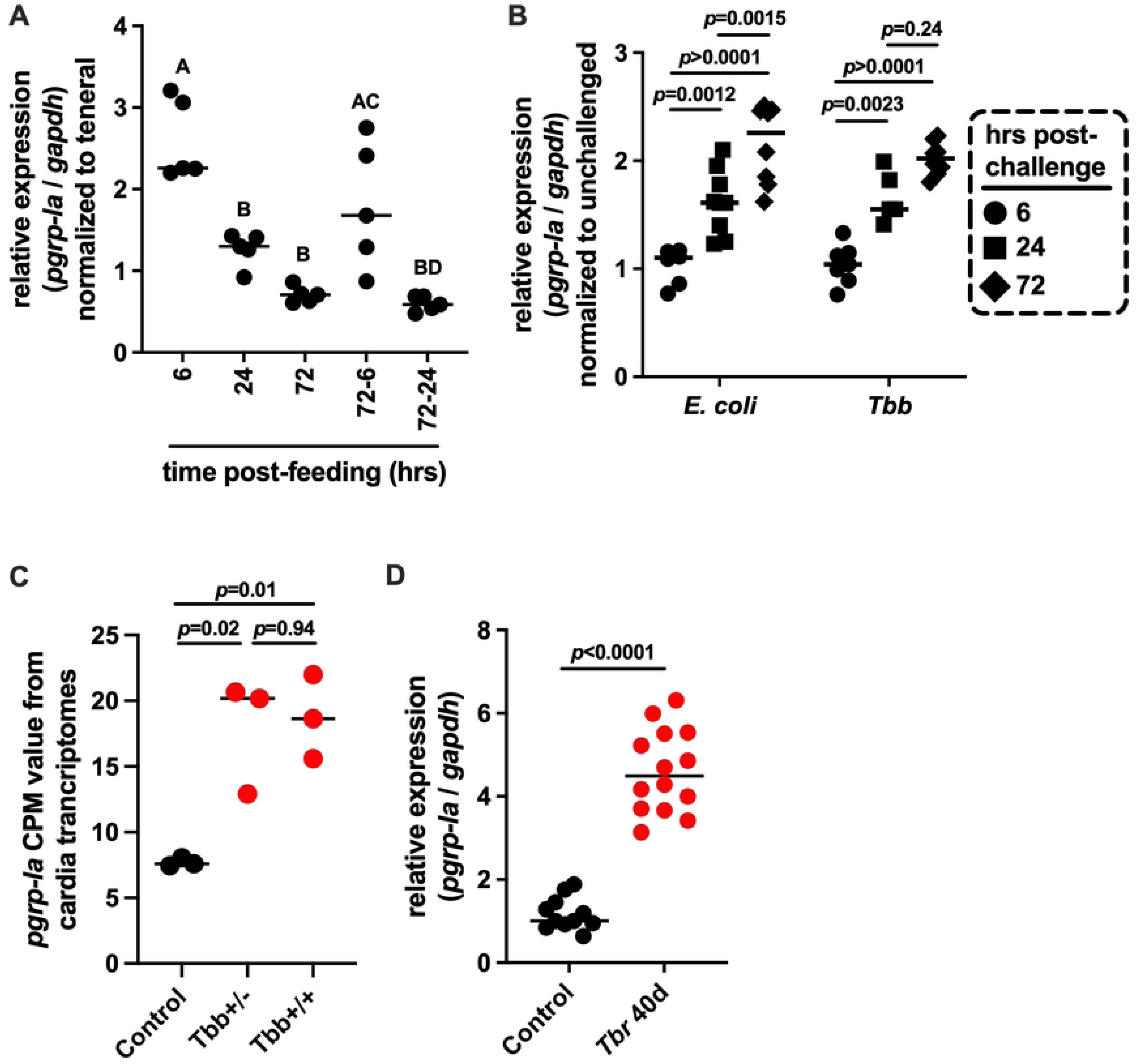
*pgrp-la* expression in response to microbial and parasitic challenge. (A) Relative *pgrp-la* expression in the cardia of flies analyzed 6, 24, and 72 hrs after their first bloodmeal, and at 6 and 24 hrs following a second bloodmeal provided after a 72-hr starvation period (72-6 and 72-24, respectively). Expression levels were normalized to *pgrp-la* expression in teneral flies. Data were analyzed using multiple Student’s t-tests. (B) Relative *pgrp-la* expression in the cardia of flies 6, 24, and 72 hrs after ingestion of a blood meal supplemented with 10^5^ CFU of *E. coli* per ml. Expression levels were normalized to those in flies that were fed a non-infectious bloodmeal. Data were analyzed using multiple Student’s t-tests. (C) *pgrp-la* transcript abundance in the cardia of uninfected flies (black circles) and flies with either gut-only (*Tbb*+/-) or gut and salivary gland (*Tbb*+/+) trypanosome infections as determined 40 days post-challenge (red circles). Data were obtained from previously published RNA-seq datasets [31] available under NCBI BioProject ID PRJNA358388. Data were analyzed using one-way ANOVA followed by Tukey’s HSD post hoc test to compare transcript abundance among groups. (D) Relative *pgrp-la* expression in the cardia of uninfected flies (black circles) and flies with gut-only *T. b. rhodesiense* (*Tbr*+/-) infections (open circles) determined 40 days post-challenge. Gene expression was assessed by qRT-PCR and normalized to *gapdh*. Each data point represents a biological replicate composed of pooled cardia tissues from five (panels A, B, D) or ten (panel C) individual flies. Data were analyzed using Student’s t-tests with correction for multiple comparisons. In panels A-D, median values are indicated by horizontal bars. Statistical analyses were performed using GraphPad Prism v.10.4.1.

In *Drosophila* [26] and *Anopheles* mosquitoes [27], *pgrp-la* is induced by microbial challenge and influences AMP expression, functioning as a PRR that regulates epithelial immunity. To investigate whether tsetse *pgrp-la* is similarly responsive to microbial exposure, we supplemented bloodmeals of teneral flies with either *Escherichia coli* (1×10^3^ CFU per ml) or BSF trypanosomes (5×10^6^ *Tbb* per ml) and measured *pgrp-la* expression at 6, 24, and 72 hrs post-feeding. No significant induction of *pgrp-la* was observed at 6 hrs for either treatment. However, both *E. coli* and *Tbb* treatment significantly induced *pgrp-la* expression at 24 hrs, with elevated expression persisting (and in the case of *E. coli* challenge, significantly increasing) through 72 hrs (Figure 3B). These results indicate that *pgrp-la* responds to microbial exposure with a delayed but sustained induction, which is consistent with a role in modulating gut immune responses during pathogen infections.

We reanalyzed data from our previous study [31], which profiled gene expression in the cardia of tsetse flies harboring either gut only infections (*Tbb*^+/-^) or gut and SG infections (*Tbb*^+/+^). This analysis revealed that *pgrp-la* was significantly induced in the cardia of both infected groups compared to uninfected controls (Figure 3C, *p*=0.02 for *Tbb*^+/-^ and *p*=0.01 for *Tbb*^+/+^). We validated these findings by performing infection experiments with a different parasite, *T. b. rhodesiense* (*Tbr*) (strain YTAT 1.1), which only establishes infections in tsetse’s gut and is unable to infect the fly’s SGs. As with *Tbb*, *Tbr* infected flies (*Tbr*^+/-^) had significantly higher *pgrp-la* expression in the cardia than did age-matched uninfected controls (Figure 3D).

Collectively, our results indicate that PGRP-LA contributes to gut homeostasis through its transient induction in the cardia during the natural feeding process, possibly in response to mechanical stretch or blood-derived signals. In the presence of microbial challenge, this response is prolonged, suggesting an additional role for PGRP-LA in gut immune defenses during pathogen exposure.

### PGRP-LA influences trypanosome colonization success

We next sought to investigate whether *pgrp-la* influences trypanosome colonization success by reducing its expression via RNAi. *Per os* treatment with anti-*pgrp-la* double stranded RNA (dsLA) reduced *pgrp-la* transcript abundance by 78% in fly midguts relative to control flies administered anti-*gfp* double stranded RNA (dsGFP) (Figure S2A). Provisioning a parasite infected bloodmeal supplemented with dsLA resulted in a significantly higher gut infection prevalence in the treatment groups compared to dsGFP controls when analyzed on day 40 post-challenge (*p*=0.04, Figure 4A). We also evaluated the SG infection status of these gut-infected flies and observed a statistically significant increase in the number of flies that had both gut and SG infections in the dsLA treatment groups (*p*=0.02, Figure 4B).

**Figure 4.**
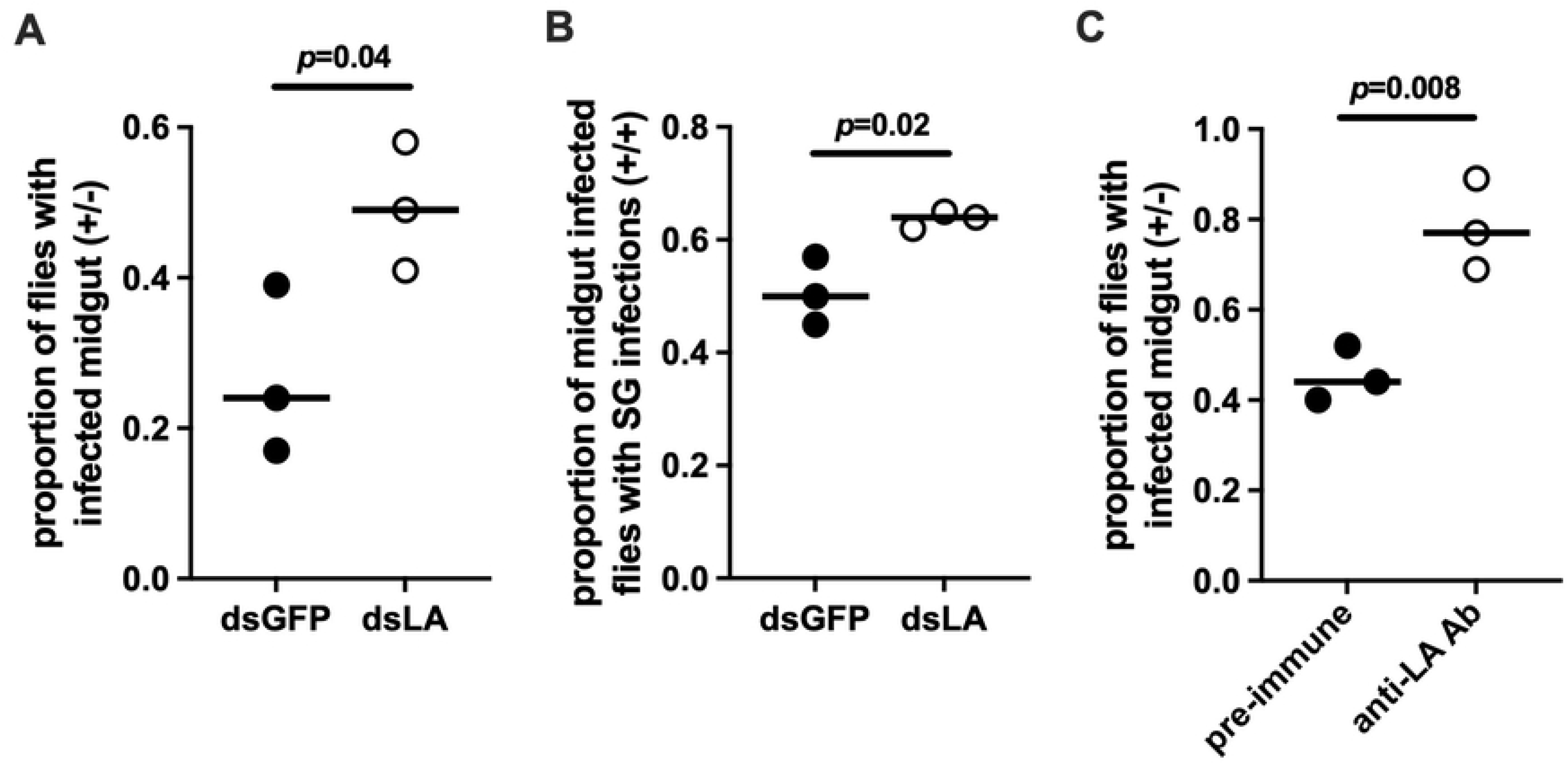
Impact of PGRP-LA on trypanosome infection outcomes. (A) Proportion of flies treated with double-stranded RNA targeting *pgrp-la* (dsLA, red circles) that developed gut-only trypanosome infections (Tbb⁺/⁻) compared to control flies treated with double-stranded RNA targeting *gfp* (dsGFP, black circles). (B) Proportion of dsLA-treated flies that developed both gut and salivary gland (SG) infections (Tbb⁺/⁺) compared to dsGFP treated controls. (C) Proportion of flies treated with anti-PGRP-LA antibodies that developed gut-only (+/-) trypanosome infections compared to flies treated with pre-immune sera. Each data point represents an individual biological replicate consisting of 25 flies. Statistical comparisons between treatment and control groups was assessed using proportion *z*-tests (GraphPad Prism v.10.4.1).

We used *E. coli* to express two recombinant proteins that encode the PGRP and non-PGRP domains of tsetse PGRP-LA, and then generated rabbit polyclonal sera against each protein. Western blot analysis using cardia tissue extracts revealed a non-specific protein band of ∼27 kDa detected with both the pre-immune and immune sera (Figure S3). In addition, immune sera generated from both recombinant proteins recognized a ∼70 kDa protein, which is about twice the expected molecular weight of the putative PGRP-LA (32 kDa). The larger than expected size observed on Western blots could result from the fact that PGRP-LA is predicted to be N-glycosylated at multiple sites, or that the protein may form a doublet *in vivo*.

We combined the two anti-LA antibodies and provided them *per os* to teneral flies three times, with the third bloodmeal also containing infectious trypanosomes (anti-LA antibody treatment group). The control group similarly received three blood meals spiked with pre-immune sera, with the third meal also containing infectious trypanosomes. Two weeks later, we monitored midgut infection prevalence microscopically and found that a significantly higher proportion of flies in the anti-LA antibody treatment group harbored trypanosomes compared to the control group; an average of 78% versus an average of 42%, respectively (*p*=0.008, Figure 4C). Collectively, our results from dsLA and anti-LA antibody treatment experiments suggest that *pgrp-la* expression levels, as well as PGRP-LA functional interactions, impact tryp infection success, possibly by influencing tsetse gut immune processes.

### Unlike in other Diptera, tsetse PGRP-LA does not interact with the fly’s Imd pathway

Barrier epithelial immune responses in *Drosophila* and *Anopheles* are regulated either by PGRP-LC alone, or through cooperative signaling with PGRP-LA. In *Drosophila*, gut specific overexpression of *pgrp-la* through transgenesis activates the expression of AMP encoding genes, including *diptericin*, *cecropin A1,* and *attacin* even in the absence of infection [24]. This AMP induction persists in *pgrp-lc* mutant flies, indicating that PGRP-LA can function independently of PGRP-LC. However, overexpression of *pgrp-la* fails to induce AMP expression in *Relish* mutant flies, indicating that PGRP-LA-mediated signaling involves the key NF-κB transcription factor downstream of Imd pathway. These findings suggest that while PGRP-LA can act as an autonomous PRR, its activity depends on the canonical Imd signaling components [24]. Unlike in *Drosophila*, in *Anopheles* mosquitoes, PGRP-LA and PGRP-LC cooperate to maintain gut epithelial homeostasis under basal non-infectious conditions [25]. Silencing either gene leads to gut dysbiosis, reduced expression of multiple immune effectors such as *attacin*, *TEP1* and *nitric oxide synthase* (NOS), and increases *Plasmodium* infection prevalence [25]. This increased susceptibility is abolished by antibiotic treatment, confirming that the anti-parasitic functions of either PRR is microbiota dependent.

Our prior studies showed that *attacin*, an Imd pathway regulated effector AMP, is strongly induced in the cardia following *E. coli* challenge through PGRP-LC signaling [23, 32]. To evaluate whether PGRP-LA modulates Imd signaling in tsetse, we reduced *pgrp-la* by feeding flies a bloodmeal supplemented with dsLA, followed by a second bloodmeal supplemented with *E. coli* (10^7^/mL of blood). This treatment achieved a 93% reduction in *pgrp-la* transcript levels (Figure S2B). Control flies received dsGFP as well as a second *E. coli* supplemented blood meal. We then quantified *attacin* expression across multiple gut-associated tissues (cardia, bacteriome, midgut, and hindgut) and found no significant differences between dsLA- and dsGFP-treatment groups (Figure 5A). These results indicate that unlike in *Drosophila* and mosquitoes, tsetse PGRP-LA does not regulate Imd-pathway activation.

**Figure 5.**
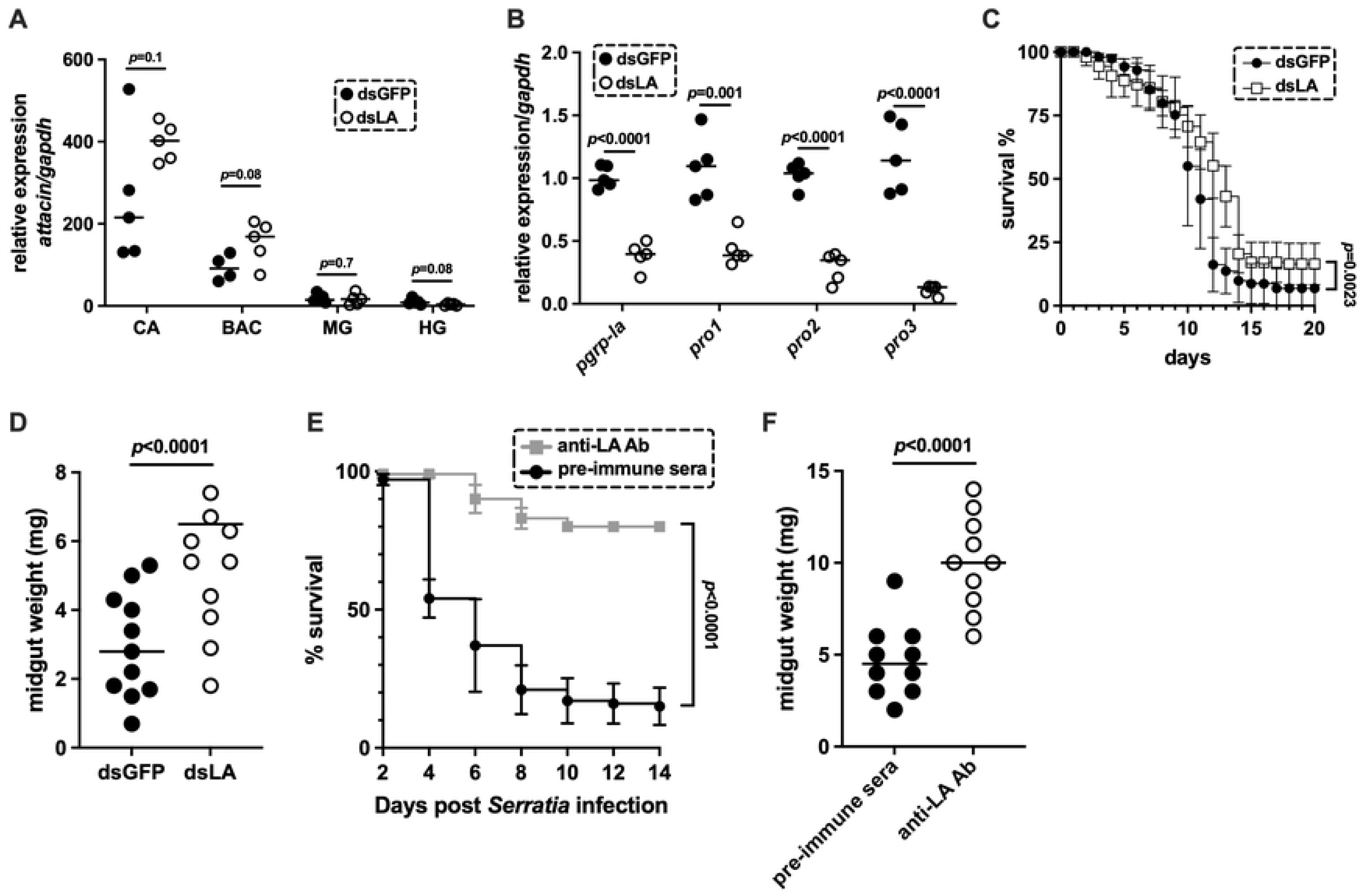
Role of PGRP-LA in regulating tsetse’s innate immune system. (A) Relative *attacin* expression in different gut compartments, cardia (CA), bacteriome (BAC), midgut (MG, and hindgut (HG) in flies treated with either dsLA or dsGFP and immune challenge with *E. coli*. (B) Relative expression of PM associated peritrophin genes (*pro1*, *pro2* and *pro3)* in cardia tissue of 14-day old flies following treatment with dsLA or dsGFP. (C) Kaplan-Meier survival curves of flies treated *per os* with dsGFP or dsLA, followed 72 hrs later by a bloodmeal supplemented with 1,000 CFU/ml of live *Serratia marcescens* (strain Db11). Flies were subsequently maintained on a normal blood diet, and mortality was monitored daily. (D) Midgut weights of flies treated with dsGFP or dsLA. Flies received dsRNAs *per os* in three consecutive blood meals, and midguts were dissected and weighed 48 hrs after the third blood meal. (E) Kaplan-Meier survival curves of tsetse treated *per os* with either pre-immune serum or anti-LA antibodies, followed 72 hrs later by a blood meal supplemented with 1,000 CFU/ml of live *S. marcescens* strain Db11. Flies were subsequently maintained on a normal blood diet, and mortality was monitored every other day. (F) Midgut weights of flies treated with either pre-immune serum or anti-LA antibodies. Flies were provided three antibody-supplemented blood meals and midguts were dissected and weighed 48 hrs after the third blood meal. For panels A and B, each data point represents a biological replicate, with each replicate containing pooled cardia tissue from five flies. Expression was normalized to the housekeeping gene *gapdh*. Data were analyzed using multiple Student’s t-tests. Survival experiments in panels C and E were conducted in two independent experiments with 25 flies per treatment per replicate, and data were analyzed using the Log-Rank (Mantel-Cox) test. In panels D and F, each data point represents one individual midgut and data were analyzed using a Student’s t-test. In panels A, B, D, and F, bars represent the median. All statistical analyses were performed using GraphPad Prism v.10.4.1.

This functional divergence may potentially reflect tsetse’s highly specialized gut microbiota, which is largely composed of maternally transmitted endosymbionts. Such a stable and beneficial microbial community may require less immune regulation, reducing the need for multiple PRRs to yield robust antimicrobial responses via the Imd-signaling. This hypothesis is consistent with the reduced complexity of the PGRP gene family in tsetse compared to *Drosophila*, which has a more diverse microbiota and an expanded set of PGRP genes.

### *pgrp-la* reduction and anti-LA antibody treatment compromise the structural integrity of tsetse’s PM

Because *pgrp-la* does not appear to coordinate epithelial immune responses in tsetse, such as the production of Imd effector protein Attacin, and is expressed preferentially in the cardia, we investigated whether it influences the formation and/or function of the tsetse PM. We found that reduction of *pgrp-la* expression levels by approximately 90% following dsRNA treatment (Figure S2C) significantly decreased the expression of multiple *peritrophin* genes that encode the PM proteins Pro1, Pro2, and Pro3 in tsetse’s cardia (Figure 5B).

To further assess the functional consequences of reduced *pgrp-la* expression on PM integrity, we conducted three complementary experiments. In the first experiment, we performed a fly microbial infection/survival assay designed to assess the structural robustness of the PM, which serves as a barrier that modulates the ability of midgut epithelial cells to immunologically detect and respond to microbial pathogens [11, 15, 31, 33]. Under normal circumstances, an intact robust PM benefits the host by preventing unnecessary immune activation by limiting epithelial exposure to gut microbiota. However, in the presence of a pathogenic microbe, this robust PM can have detrimental consequences on host survival as it may delay the detection of microbe-associated molecular patterns (MAMPs), thereby postponing the initiation of epithelial immune responses. Such a delay would allow pathogenic microbes to replicate unchecked, ultimately resulting in host death due to systemic infection. To test this, flies from both the dsLA treatment and dsGFP control groups were given a bloodmeal supplemented with 1×10^3^ CFU of entomopathogenic *Serratia marcescens* strain db11. We observed that dsLA treated individuals, with reduced *pgrp-la* expression, survived significantly longer than did dsGFP controls (Figure 5C). This extended survival suggests that the PM in dsLA-treated flies was structurally compromised, allowing earlier immune detection and response to the invading pathogen. In contrast, the intact PM in control flies likely delayed immune activation, contributing to increased susceptibility and earlier mortality.

In the second experiment, we quantified the correlation between PM functionality and midgut weight, which serves as a reflection of digestive homeostasis [11, 15, 31, 33]. When we weighed midguts excised from both the dsLA treatment and dsGFP control groups 48 hrs after their last blood meal, we found that midguts from the dsLA treatment group weighed significantly more than those from the control dsGFP group (Figure 5D, p*<*0.0001). This finding further indicates a disruption in PM integrity and digestive functionality as a consequence of PGRP-LA reduction.

Finally, we provided two blood meals supplemented with anti-LA antibodies and a third blood meal supplemented with both anti-LA antibodies and entomopathogenic *Serratia*. A similar control group was established using pre-immune sera. This experimental design was used to assess the impact of anti-LA antibody treatment on fly survival. Flies treated with anti-LA antibodies survived significantly longer following the *Serratia* challenge than did control group (Figure 5E). Additionally, antibody-supplemented treatment flies presented with significantly heavier midguts compared to control flies (Figure 5F).

Taken together, these results indicate that the PGRP-LA protein functions in pathway(s) that regulate the expression of proteins essential for the structural integrity of the tsetse gut PM barrier. As such, PGRP-LA has the potential to influence both the outcome of infections with pathogenic bacteria and the transmission success of parasitic trypanosomes.

### sVSG and miR-275 regulate pgrp-la expression

We next sought to investigate whether PGRP-LA may function as a receptor or signaling molecule in pathways regulating PM synthesis, which is critical for gut homeostasis and for susceptibility to trypanosome infection. In previous studies, we demonstrated that supplementing tsetse’s blood diet with BSF trypanosomes, BSF extracts, or purified BSF sVSGs, transiently reduces expression of *peritrophin* genes in the cardia, resulting in compromised PM integrity [9]. To determine whether *pgrp-la* expression in the cardia is similarly affected by trypanosome-derived components, we fed teneral flies blood supplemented with either purified sVSG (treatment) or BSA (control) and measured *pgrp-la* expression 48 hrs post-feeding. We found that sVSG exposure significantly decreased *pgrp-la* expression in the cardia (*p*=0.046, Figure 6A), indicating that mammalian trypanosome-derived molecules that impair PM structure also downregulate *pgrp-la*.

**Figure 6.**
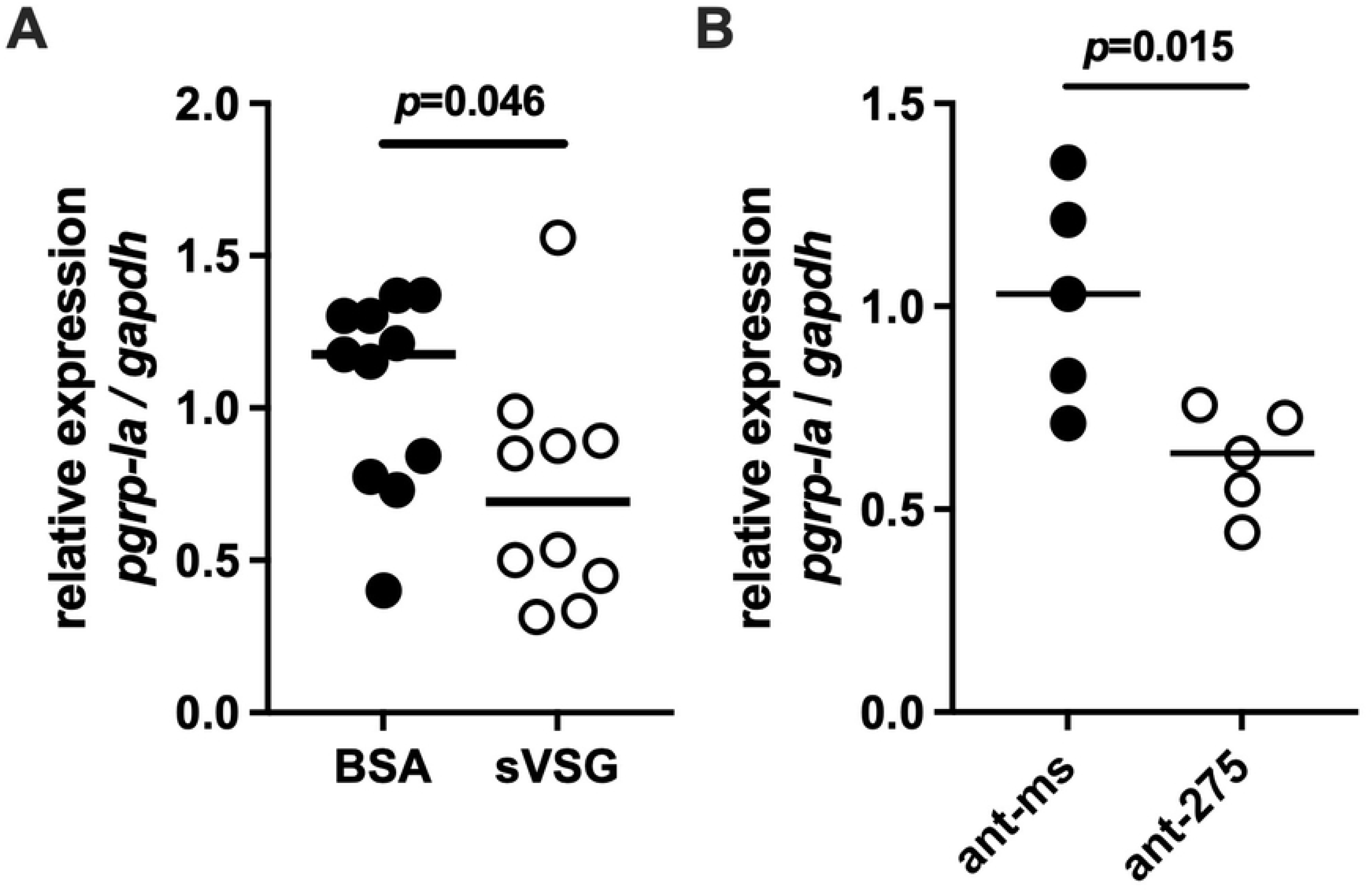
Regulation of *pgrp-la* expression in the tsetse cardia. (A) Relative expression of *pgrp-la* in the cardia tissue of 11-day old flies fed blood meals supplemented with either bovine serum albumin (BSA, black circles) or soluble variant surface glycoprotein (sVSG, open circles). (B) Relative expression of *pgrp-la* in the cardia tissue of 11-day old flies fed blood meals supplemented with either an antagomir targeting mir-275 (ant-275, black circles) or a scrambled control antagomir (ant-ms, open circles). In both panels, each data point represents a biological replicate composed of three pooled cardia collected 72 hrs after supplementation. Bars indicate the median. Quantitative real-time PCR (qRT-PCR) results were normalized to expression of the housekeeping gene *gapdh*. Data were analyzed using Student’s t-tests (GraphPad Prism v.10.4.1).

We previously showed that sVSG uptake by cardia cells reduces the expression of *microRNA-275* (*miR-275*), which regulates PM-associated *peritrophin* genes expression in the cardia and consequently PM integrity [15]. A similar decrease in *peritrophin* gene expression was observed following depletion of *miR-275* levels via synthetic anti-*miR-275* antagomirs [15]. To determine whether *pgrp-la* expression is also regulated by *miR-275*, we treated flies *per os* with anti-*miR-275* antagomirs (treatment) and measured *pgrp-la* transcript levels. Flies treated with the antagomir exhibited a significant reduction in *pgrp-la* expression compared to those receiving a missense control antagomir (*p*=0.015; Figure 6B). These results suggest that *pgrp-la* may act as a sensor that participates in the *miR-275*–regulated pathway controlling PM synthesis in the cardia following blood meal ingestion.

Together, these findings highlight a regulatory cascade in which BSF trypanosome surface antigens (sVSGs) suppress *miR-275* expression, leading to downstream reductions in both *peritrophin* and *pgrp-la* expression, thus ultimately weakening the PM barrier. While it remains unclear how trypanosome-derived components mechanistically interact with the PGRP-LA receptor, and how *miR-275* levels are modulated by their yet to be determined target proteins, our data suggest that this regulatory network is central to controlling PM integrity and host susceptibility to infection. Future studies are needed to obtain biochemical evidence that PGRP-LA physically binds sVSG. Ligand-binding assays, such as co-immunoprecipitation or surface plasmon resonance can help test this hypothesis.

## Discussion

Tsetse flies exhibit a markedly reduced repertoire of PGRP-encoding genes. Specifically, their genomes encode only four to six members of the PGRP family [17] compared to the 13 and 17 found in *Drosophila melanogaster* and *Musca domestica*, respectively [34]. With so few PGRPs across the *Glossina* genus in comparison to other Dipteran insects, the retention of a structurally conserved PGRP-LA, complete with key domains such as the RHIM-like motif, strongly suggests that this receptor plays an essential and non-redundant immune-related function in tsetse. Our data indicate that, despite this structural conservation, tsetse PGRP-LA has diverged functionally. Unlike its *Drosophila* and mosquito orthologues, which regulate Imd signaling and AMP production during microbial challenge, tsetse PGRP-LA appears to primarily regulate the function of a different immune barrier, the midgut-associated PM.

Tsetse *pgrp-la* transcripts are enriched in gut-associated tissues, with the highest expression detected in the PM-secreting cardia located at the junction of the foregut and midgut. This tissue-specific enrichment is different from expression patterns in *Drosophila* where *pgrp-la* is enriched in barrier epithelia, including the gut and tracheae [26]. In tsetse, *pgrp-la* transcript abundance increases shortly after blood feeding and remains elevated following *per os* exposure to either bacteria or trypanosomes. *Pgrp-la* is also highly expressed in flies that house mature parasite infections, relative to age-matched uninfected controls. In *Anopheles* mosquitoes, *pgrp-la* expression in the midgut also increases following a blood meal, likely in response to microbiota proliferation, as this induction is abolished in axenic mosquitoes that lack their gut microbiota [27]. The enteric microbiota of field captured, and laboratory reared tsetse flies is highly taxonomically streamlined compared to that of mosquitoes and other arthropod vectors [35]. In fact, the laboratory reared flies used in this study house only facultative *Sodalis* in their gut lumen as the obligate *Wigglesworthia* is confined within bacteriocytes that compose tsetse’s bacteriome, which is attached to the hemocoel-exposed surface of the fly’s gut [36]. Given the small population size of *Sodalis* in tsetse’s gut [37] and the very early timing of the *pgrp-la* upregulation we noted post blood-feeding (6-h post feeding), we speculate that induction of the gene in tsetse following consumption of bloodmeal may be driven by stretch of the cardia and/or a molecular component of the blood. The outcome would represent a significant evolutionary divergence of *pgrp-la* biology in tsetse compared to mosquitoes.

Reduction of *pgrp-la* expression in tsetse via RNAi led to a marked increase in the proportion of flies with midgut trypanosome infections compared to controls. A similar increase was observed when infectious bloodmeals were supplemented with anti-PGRP-LA antibodies, implicating protein level interference with PGRP-LA function. Importantly, antibody-mediated blocking of PGRP-LA reproduced the RNAi phenotypes, suggesting that PGRP-LA function extends beyond transcriptional regulation and may involve protein-level sensing or signaling. To explore how PGRP-LA contributes to parasite resistance, we examined its role in two major pathways known to affect tsetse vector competence: immune effector responses and the structural integrity of the PM physical barrier. Knockdown of *pgrp-la* did not enhance expression of *attacin*, an Imd-regulated antimicrobial peptide with trypanolytic activity [38], suggesting that PGRP-LA does not function through canonical immune signaling. Instead, *pgrp-la* depletion significantly reduced expression of several *peritrophin* genes and induces PM dysfunction. Moreover, both RNAi- and antibody-treated flies exhibited phenotypes consistent with PM dysfunction, including decreased susceptibility to entomopathogenic *Serratia* and increased midgut weight following blood feeding, indicative of impaired digestion. These phenotypes closely mirror those reported following RNAi-mediated knockdown of *chitin synthase* or *peritrophin* genes [11], and mimic the effects of microRNA-275 suppression, which similarly weakens PM integrity and increases midgut infection prevalence [15]. Taken together, these results incriminate PGRP-LA as a key regulator of gut homeostasis, acting to maintain PM barrier structural and functional integrity, as opposed to stimulating epithelial immune responses. Future studies are needed to define downstream signaling pathways through which PGRP-LA modulates *peritrophin* expression and its connections to the miR-275/Wingless regulatory axis or other gut epithelial responses. Importantly, antibody feeding studies may have off-target or indirect effects that complicate interpretation of protein-level function. Future work will also need to address these limitations through direct ligand interaction assays and pathway dissection.

Our findings highlight tsetse’s cardia as a critical anatomical and functional site of vector-microbe interactions. Tsetse PGRP-LA is most abundantly expression in the cardia that constitutively synthesizes Type II PM that lines tsetse’s midgut. In addition, the cardia is also an active immune niche where oxidative and antimicrobial defenses are produced [11, 23]. Spatial confinement of PGRP-LA to the cardia suggests that it acts as a localized infection sensor, integrating signals from ingested microbes or parasite-derived factors to regulate PM formation. Such a regulatory role is supported by significantly elevated *pgrp-la* expression in the cardia of flies with mature trypanosome infections, indicating that PGRP-LA may be upregulated to restore or maintain PM function. However, this defense mechanism may also be manipulated by trypanosomes. Specifically, VSGs shed from newly acquired BSF trypanosomes are internalized by tsetse’s cardia where they suppress the expression of *miR-275*. This process transiently reduces the expression of PM-associated genes and results in disruption of the barrier [15] and provides a temporary but critical window of vulnerability that facilitates parasite migration across the PM and subsequent midgut colonization. Interestingly, PGRP-LA knockdown or antibody feeding produces similar outcomes, suggesting that trypanosomes may target both *miR-275* and PGRP-LA expression to enhance early colonization success. Whether sVSGs directly interact with PGRP-LA or act through downstream intermediates remains unknown. As a PRR, PGRP-LA may also interact with unidentified parasite-derived ligands or host factors that modulate its stability or putative signaling activity. Our results support a model in which the cardia acts as the “gatekeeper” of the tsetse midgut where it physically detects passage of blood and microbes and in response regulates PM formation and host defenses. PGRP-LA has a critical function for pathogen sensing and gut barrier formation, effectively regulating the downstream pathways that maintain gut homeostasis and immune surveillance. As such PGRP-LA provides a potential target for interventions that block trypanosome transmission through PM strengthening.

## Conclusions

Our study uncovers a previously uncharacterized role for the PPR PGRP-LA in maintaining the integrity of the tsetse fly PM, a critical barrier to early trypanosome infection success. Given that PM disruption facilitates parasite transmission, bolstering PGRP-LA function may represent a promising strategy to reduce tsetse vector competence. Understanding this natural defense mechanism could inform the development of novel interventions targeting PGRP-LA or its downstream effectors in tsetse and other disease-transmitting insects, thus contributing to broader efforts to control trypanosomiasis and related vector-borne diseases. Future work can now focus on elucidating the downstream signaling cascades that link PGRP-LA to Peritrophin gene expression and on identifying how trypanosome molecules interact with PGRP-LA and interfere with this regulatory pathway. From an applied perspective, such knowledge could reveal novel targets for blocking transmission - for example, interventions that block VSG’s effects or boost PGRP-LA activity to reinforce PM integrity.

## Materials and Methods

### Tsetse flies, bacteria, and trypanosomes

Tsetse flies (*Glossina morsitans morsitans*) used in this study were maintained in the Yale School of Public Health insectary at 25°C with 65% relative humidity. Flies received defibrinated bovine blood (Lampire Biologicals) every 48 hrs using an artificial membrane feeding system. Flies referred to as ‘teneral’ were unfed adults newly eclosed from their pupal case (≤ 48h).

Bloodstream form *Trypanosoma brucei brucei* strain RUMP 503 (*Tbb*) or *T. b. rhodesiense* YTat 1.1 (*Tbr*) were expanded in rats and aliquots of infectious blood were stored at -80°C. Flies were infected by supplementing the first blood meal of teneral flies with 1×10^6^ parasites/ml. All experiments were performed in accordance with Yale IACUC protocol number 2023-07266.

For survival assays, *Serratia marcescens* strain Db11 was grown overnight in LB medium at 27°C. Prior to supplementation with *Serratia*, the blood was heat inactivated at 56°C for 1 h.

### Analysis of *pgrp-la* and *PGRP-SA* expression in response to trypanosome infection

Teneral female flies were given a blood meal supplemented with 5×10^6^ BSF *Trypanosoma brucei brucei* (*Tbb*) per ml of blood, and thereafter fed normal blood every other day. Fourteen days later (and 48 hrs after their last bloodmeal) cardia, bacteriomes, and post-bacteriome midguts were collected and stored at -80°C for later processing and analysis of *pgrp-la* (VectorBase gene ID, GMOY006093) and *PGRP-SA* (GMOY009549) expression (see ‘RNA extraction, cDNA synthesis and qRT-PCR’ subsection below).

### Spatial analysis of *pgrp-la* expression and comparison to the *Drosophila* ortholog

To analyze *pgrp-la* expression in different compartments of tsetse’s whole gut, 10-day old adult female flies were microscopically dissected 48h after the last bloodmeal. Either whole guts, or cardia, bacteriomes, post-bacteriome midguts, and hindguts, were dissected and stored at -80°C for later processing and analysis of *pgrp-la* expression (see ‘RNA extraction, cDNA synthesis and qRT-PCR’ subsection below). Twelve and six biological replicates were generated for whole digestive track and each dissected part, respectively. Expression of *pgrp-la* was evaluated using quantitative real-time PCR (qRT- PCR) and normalized to the housekeeping gene *gadph* for each sample. The resulting average expression of each part of the digestive track was then normalized to the average expression of the whole digestive track. For comparative analysis, *Drosophila pgrp-la* expression values corresponding to the different regions of the digestive track were obtained from the FlyGut database, (https://flygut.epfl.ch/expressions/show_by_gene?synonym=CG32042) [39].

### Temporal analysis of *pgrp-la* expression in response to microbial challenge and infection

To evaluate the timing of *pgrp-la* expression in response to microbial challenge, teneral female flies were given a blood meal supplemented with either 1×10^5^ CFU of *Escherichia coli* or 5×10^6^ BSF *Trypanosoma brucei brucei* (*Tbb*) per ml of blood. Cardia tissues were dissected at either 6, 24, or 72 hrs post-challenge. Control groups consisted of age-matched unchallenged flies.

*pgrp-la* expression patterns in cardia of 40-day old midgut only (*Tbb*^+/-^) and midgut and SG (*Tbb*^+/+^) *Tbb* infected tsetse flies were determined by mining RNA-seq data obtained in our previous study [31]. Relevant RNA-seq read data files can be accessed via NCBI BioProject ID# PRJNA358388. To evaluate cardia *pgrp-la* expression in response to *Tbr* infection, teneral females were offered a blood meal containing 5×10^6^ BSF *Tbr* per ml of blood. Forty days post-challenge (and three days after their last blood meal) flies were dissected and their midguts were microscopically evaluated to determine *Tbr* infection status. Cardia from midgut infected flies (*Tbr*^+/-^), and cardia from age matched unchallenged controls, were collected and stored at -80°C for later processing and analysis (see ‘RNA extraction, cDNA synthesis and qRT-PCR’ subsection below).

### pgrp-la RNAi

PGRP-LA and green fluorescent protein specific dsRNAs (dsLA and dsGFP, respectively) were synthesized using a MEGAscript T7 *In Vitro* Transcription kit according to the manufacturer’s (Invitrogen) protocol. All dsRNA PCR primer sequences (Table S1) were confirmed against tsetse genomic scaffolds and RNA-seq libraries (www.vectorbase.org) to eliminate possible off target effects. In addition, the PCR products were sequenced before dsRNA synthesis to confirm that they encoded the target genes.

### *pgrp-la* knockdown and effects on host physiology

*Trypanosome infections:* Seventy-five teneral female flies received a blood meal containing 5×10^6^ BSF *Tbb* per ml of blood and 8µg of either dsLA or dsGFP in 20μl blood. After forty days, parasite infection prevalence was microscopically determined in the gut and SG of all flies.

#### Antimicrobial protein and PM associated gene expression

Eight-day old adult flies received 8µg of dsLA (treatment group) or dsGFP (control group) per 20μl blood (the approximate volume a tsetse fly imbibes each time it feeds) in two consecutive blood meals. To measure the impact of *pgrp-la* knockdown on *attacin* expression, flies were provided live *E. coli* DH5ɑ (10^7^/ml of blood) in their second dsRNA containing blood meal. Twelve hrs later fly guts were microscopically dissected, divided into cardia, bacteriome, post-bacteriome midgut, and hindgut sections, and stored at -80°C for later processing and analysis of *attacin* expression (see ‘RNA extraction, cDNA synthesis and qRT-PCR’ subsection below). To measure the impact of *pgrp-la* knockdown on the expression of PM associated genes, cardia tissues were dissected from flies 12 hrs after receiving their second dsRNA containing blood meal (no *E. coli* were used for this experiment) and stored at -80°C for later processing and analysis of *pro1, pro2, and pro3* expression.

#### Host survival following challenge with an entomopathogenic bacterium

To evaluate host survival as a consequence of PM integrity, we used *Serratia* infection assays, following the methods described in [11, 15]. Eight-day old adult flies received 8µg of dsLA (treatment group) or dsGFP (control group) per 20μl blood in their first two consecutive blood meals. Forty-eight hrs after receiving dsRNAs, all flies were given another blood meal supplemented with 10^3^ CFU/ml of *S. marcescens* strain Db11. Thereafter, all flies were maintained on a blood diet supplemented with their respective dsRNAs, and mortality was recorded daily for 14 days.

#### Midgut weight assay

To evaluate the impact of dsLA-treatment on midgut weights, we provided 8-day old flies with blood meals supplemented with 8 µg/20 µl of either dsGFP or dsLA. Twenty-four hrs after the dsRNA supplemented bloodmeal, fly guts (cardia to the midgut/hindgut junction) were dissected and weighed to the nearest 0.1 mg.

### Anti-PGRP-LA antibody production

Two polyclonal antibodies were developed in rabbits against distinct PGRP domains of tsetse PGRP-LA. The first antibody (anti-LA1) targets the domain encoded by base pairs 415 to 853 of the putative mature *pgrp-la* transcript, while the second antibody (anti-LA2) targets the domain encoded by base pairs 763 to 1011 of the same transcript. The anti-LA1 domain was PCR-amplified and cloned into the pET-28a vector and transformed into BL21 cells. Expression was induced by adding IPTG to a final concentration of 1.0 mM when cells reached OD_600_ 0.6-1.0. Cells were harvested by centrifugation at 12,000 rpm for 15 min at 4°C, and the resulting pellet was resuspended in lysis buffer (Merck, 71456-3). Inclusion bodies were collected by centrifugation at 12,000 rpm for 20 min at 4°C and re-suspended in inclusion buffer (50mM Tris, 100mM NaCl, 8M Urea, 5% Glycerol, 1mM PMSF, 10mM DTT). The anti-LA2 domain was PCR-amplified and cloned into the pAE vector [40] and the rec-Protein was similarly expressed the BL21 cells. Two mg of each rec-protein was analyzed by SDS-PAGE, and the protein bands were excised and sent to Cocalico Biologicals for antibody generation in two rabbits for each antibody.

Western blot analysis was performed using pooled cardia from 5 individual flies homogenized in 100 μl of lysis buffer (8 M urea, 2% SDS, 5% β-mercaptoethanol, 125 mM tris-HCl), centrifuged (12,000 rpm) at 4°C for 10 min, and heated at 95°C for 5 min. Twenty-five microliters of supernatant was combined with 4x Laemmli sample buffer, run on a 4-15% SDS-PAGE gel (Bio-Rad 456-1085), transferred to nitrocellulose membranes, and blocked 3% BSA, 0.1% tween 20, in PBS) for 1 hr. Blots were incubated overnight at 4°C with either pre-immune sera or with either anti-LA1 or anti-LA2 antibodies (both diluted at 1:10,000 in blocking buffer), exposed to HRP-conjugated anti-rabbit secondary antibody (diluted at 1:20,000 in blocking buffer) for 1 hr, and visualized using a SuperSignal West Pico Chemiluminescent Substrate kit according to the manufacturer’s (Thermo Scientific 34577) protocol Figure S3.

### Anti-LA antibody effects on host physiology

#### Trypanosome infection assay

Seventy-five teneral female flies received a blood meal containing either both anti-LA antibodies or pre-immune serum alone (constituting no more than 15% of the blood mixture) and 5×10^6^ BSF *Tbb* per ml of blood. All groups received two additional blood meals supplemented with either anti-LA antibodies or pre-immune sera and were then maintained on a normal blood diet for 14 days. Guts were dissected and trypanosome infection status was microscopically determined.

#### Midgut weight assay

To evaluate the impact of feeding flies anti-LA antibodies on midgut weight, we provided 8-day old flies with blood meals supplemented with either both anti-LA antibodies or with pre-immune serum alone (constituting no more than 15% of the blood mixture). Twenty-four hrs after antibody *per os* supplementation, fly guts (cardia to the midgut/hindgut junction) were dissected and weighed to the nearest 0.1 mg.

##### Expression of *pgrp-la* after sVSG ingestion

Soluble variant surface glycoproteins (sVSG) were prepared by the method described in [15]. To assess the effect of sVSG, either purified sVSG (treatment, 1µg/mL) or bovine serum albumin (BSA) (control, 1µg/mL) provided to 8-day old flies in a blood meal. Seventy-two hrs post treatment, cardia tissues were collected and stored at -80°C for later processing and analysis of *pgrp-la* expression (see ‘RNA extraction, cDNA synthesis and qRT-PCR’ subsection below).

##### Expression of *pgrp-la* following antagomir miR-275 treatment

An antagomir targeting tsetse mir-275 (treatment, ant-275) and a randomly scrambled “missense” antagomir (control, ant-ms) were synthesized at the Yale University Keck Center (Oligo Synthesis Resource). Antagomirs were designed and used as previously described [15, 41]. Briefly, antagomirs were resuspended in sterile water (200 μM solution) and provided to 8-day old flies in the blood meal (100 μl per ml of blood). Twenty-four hrs post treatment cardia tissues were collected and stored at -80°C for later processing and analysis of *pgrp-la* expression (see ‘RNA extraction, cDNA synthesis and qRT-PCR’ subsection below).

### RNA extraction, cDNA synthesis and qRT-PCR

All RNAs were extracted from dissected tissue samples using the Direct-zol RNA Miniprep Kit (Zymo Research, Cat ZR2050) following the manufacturer’s protocol. One hundred nanograms of RNA was then reverse transcribed into cDNA using the iScript cDNA synthesis kit (BioRad), and cDNA was subjected to RT-qPCR analysis (two technical replicates were used for each sample). Relative expression (RE) was calculated using the formula RE = 2-ΔΔCt, and normalization was performed using the constitutively expressed *Glossina gapdh* gene. All PCR primers used in this study are listed in Table S1. RT-qPCR was performed on a T1000 PCR detection system (Bio-Rad, Hercules, CA) under the following conditions: 8 min at 95°C; 40 cycles of 15 s at 95 °C, 30 s at 57 °C or 55 °C, 30 s at 72 °C; 1 min at 95 °C; 1 min at 55 °C and 30 s from 55 °C to 95 °C. Each 10 μl reaction contained 5 μl of iTaq^TM^ Universal SYBR® Green Supermix (Bio-Rad), 1 μl cDNA, 2 μl primer pair mix (10 μM), and 2 μl nuclease-free H_2_O.

### Sample sizes and statistical analyses

Sample sizes are provided in the corresponding figure legends or indicated graphically as individual data points in dot plots. Biological replication refers to the use of distinct groups of flies treated in separate, independently conducted experimental procedures.

All statistical analyses were performed using Prism v.10.4.1 (GraphPad software). Relative expression of *pgrp-sa* and *pgrp-la* (Figure 1) and the effects of dsRNA treatment on *attacin* levels (Figure 5) were analyzed using two-way ANOVA, with gene expression as the dependent variable and tissue and infection status as independent variables. When significant effects were detected, pairwise comparisons were performed using post hoc Tukey HSD tests.

Relative expression of *pgrp-la* over time post-feeding in control, *E. coli*-infected and *Tbb-*infected flies, *pgrp-la* CPM, and relative expression of *pgrp-la* in *Tbr*-infected flies (Figure 3), as well as gene expression following dsLA treatment and midgut weights (Figure 5) were analyzed using Student’s *t*-tests with correction for multiple comparisons when appropriate. Relative expression of *pgrp-la* following sVSG ingestion and ant-275 treatment (Figure 6) were also evaluated using Student’s *t*-tests. Pairwise comparisons of parasite infection proportions were conducted using proportion *Z*-tests (Figure 4), and survival curves were compared using Log-Rank tests (Figure 5)

## Data availability

All relevant data are within the manuscript and its Supporting Information files. RNA-seq datasets are available at NCBI under accession number PRJNA358388.

## Acknowledgements

We are thankful for tsetse materials provided from the United Nations International Atomic Energy Association Coordinated Research Project titled Improvement of Colony Management in Insect Mass-Rearing for SIT Applications (to SA and BW). We thank Mr. Sidiya Mbodj for technical support with the tsetse husbandry and Aksoy laboratory members for their critical comments on the manuscript.

## Supplemental figure legends

**Figure S1: Characterization of *pgrp-la* genomic locus and protein sequence.**

(A) Schematic view of *pgrp-la* and its putative product. The putative product contains an extracellular PGRP domain (green), a transmembrane domain (TM, dark purple) and a cytoplasmic region with a RHIM-like domain (blue). Coding sequence (CDS) regions lacking known functional domains are shown in gray. No signal peptide was predicted. The regions used to generate the recombinant proteins for antibody production (anti-LA1 and anti-LA2) are also indicated.

(B) Protein sequence alignment of the RHIM-like domain from *Glossina morsitans* (Gm) PGRP-LC and PGRP-LA, and *Drosophila melanogaster* (Dm) PGRP-LC and PGRP-LA. Residues marked with red asterisk are essential for DmPGRP-LC RHIM-like domain to activate IMD pathway, while black asterisk are contributing to the activation without being mandatory. The grey shaded amino acids depict conserved or similar residues written in black and white, respectively.

(C) Protein sequence alignment of the PGRP domain of PGRP-LA from *G. morsitans morsitans* (Gm), *G. pallidipes* (Gp), *G. austeni* (Ga), *G. fuscipes fuscipes* (Gf), *G. p. palpalis* (Gpp), *Musca domestica* (Md), *Stomoxys calcitrans* (Sc), *Ceratitis capitata* (Cc), *Aedes aegypti* (Aa) and *D. melanogaster* (Dm). PGRP domain from DmPGRP-LC, for which the crystal structure has been resolved, is presented as a reference. Residues interacting with bacterial PGN in DmPGRP-LC are underlined at the bottom of the alignment. Asterisks indicate amino acids required for amidase activity (numbering starts at the beginning of the PGRP domain). Alpha helices (α) and Beta strands (β) are represented in blue and pink, respectively. Grey background displays conserved amino acids with identities and similarities written in black and white, respectively. Red shaded residues indicate *Glossina* specific substitution. Yellow shading indicates residues conserved in *Glossina* plus another taxa.

**Fig S2. Efficiency of dsRNA-based gene knockdown**

Relative *pgrp-la* expression in cardia of dsLA (treatment) and dsGFP (control) treated flies prior to challenge with (A) trypanosomes, (B) *E. coli*, and (C) *Serratia*. Each dot on the graph represents one biological replicate, each replicate containing cardia from five dsRNA treated individuals. Statistical significance was determined via student’s t-test (GraphPad Prism v.10.4.1).

**Fig S3. Western blot analysis of anti-LA1 and anti-LA2 antibodies**

Western blot analysis was performed using protein extracts from cardia of 5 individual flies. Blots were incubated overnight at 4°C with either pre-immune sera or with either anti-LA1 or anti-LA2 antibodies (both diluted at 1:10,000), exposed to HRP-conjugated anti-rabbit secondary antibody (diluted at 1:20,000) and visualized using a SuperSignal West Pico Chemiluminescent Substrate kit. The protein size standards are indicated.

